# Open-Source Platform for Kinematic Analysis of Mouse Forelimb Movement

**DOI:** 10.1101/2024.01.31.578239

**Authors:** Daniil Berezhnoi, Hiba Douja Chehade, Hong-Yuan Chu

## Abstract

We present an open-source behavioral platform and software solution for studying fine motor skills in mice performing reach-to-grasp task. The behavioral platform uses readily available and 3D-printed components and was designed to be affordable and universally reproducible. The protocol describes how to assemble the box, train mice to perform the task and process the video with the custom software pipeline to analyze forepaw kinematics. All the schematics, 3D models, code and assembly instructions are provided in the open GitHub repository.

**Graphical abstract:** 

## Before you begin

Acquisition and execution of complex and skilled motor activity involve synergistic interaction of the cerebral cortex, the basal ganglia, the cerebellum and the spinal cord. The reach-to-grasp task in rodents has been established and used for decades to investigate neurobiological mechanisms underlying the skilled motor activity and its impairments in models of human diseases (Whishaw and Pellis, 1990; Miklyaeva et al., 1994; Whishaw, 2000; Xu et al., 2009; Azim et al., 2014; Guo et al., 2015a; Guo et al., 2015b; Bova et al., 2020; Aeed et al., 2021; Calame et al., 2023). In the present work, we describe the design and fabrication of the hardware-software platform for mouse reach-to-grasp task and use of this platform for mice training and recording behavioral videos. As a part of the platform, we present an open-source software for video analysis and kinematic quantification of mouse fine motor skills. We share our design and analysis pipeline to promote the reproducibility and availability of this protocol to the wide open-science community. All the schematics, code, 3D models, printing and assembly instructions are provided in the open GitHub repository (https://github.com/BerezhnoyD/Reaching_Task_VAI; doi/10.5281/zenodo.7383917). The reaching box is programmed with Arduino IDE and can be used as a device for automated behavioral training and data acquisition. In addition, it has the features to log basic behavioral data (touch/beam sensors) and trigger or synchronize external devices for electrophysiology recording, *in vivo* imaging, optogenetic stimulation etc. Last, we trained a cohort of wildtype C57BL6J mice using this platform and performed kinematic analysis of forelimb movement. We found that most animals successfully acquired the forelimb reaching skills. Troubleshooting tips that we found useful in our practice were reported, which can be beneficial to researchers entering this field.

### Institutional permissions

All animal procedures in this study were reviewed and approved by the Van Andel Institute Animal Care and Use Committee (IACUC; Protocol# 22-02-006).

### Behavioral box construction

#### Timing: 2-3 days

##### Steps for manufacturing parts

1. Design of 3D parts and Plexiglas sheets and assembly of the box. There are 4 main components that you will need to assemble the behavioral box (See Fig.1B for the overall schematics of the box and Fig.1A,C,D for the view from different angles, Key Resource table):
  - Plexiglas sheets cut to size and drilled,
  - 3D printed parts, such as corners holding the sheets together,
  - small screws and nuts kit, and
  - metal rods for the floor.

**Fig.1.** Overview of the 3d printed reaching box from (A) the front, (B) orthogonal view, (C) the top, (D) the side, showing the main features of the behavioral apparatus: 1) rotating disk feeder for automatic food delivery not obstructing the view of the mouse behavior in the box, 2) vertical slit for the animal to reach through, 3) the mirror system allowing to record reach-to-grasp movement from multiple views and reconstruct the trajectory, 4) metal grid floor with frontal bars connected to the controller and used as touch sensors, 5) autonomous experiment controller allowing the full control of the experiment and recording of the animal behavior in the box along with providing the synchro-signal for ephys recording or other experimental devices, 6)single high-speed camera for capturing behavior.
2. Assembly instructions for the training box.
  1. See the Fig. 2 for the dimensions of each Plexiglas part.

**Fig.2.** Blueprints of the reaching box that can be used to manufacture it using the Plexiglas sheets and 3d printed parts. A – side wall with the holes for the metal floor and the screws (3mm Plexiglas), B – front wall with a slit for animal paws (0.5 mm Plexiglas), C – models for 3d printed parts of the box, D – back wall with the holes for the screws (3mm Plexiglas). All the parts on the pictures A,B,C are shown with the same scale, the scale on the D is different. All the blueprints with precise dimensions and 3d models are provided in Supplementary materials (Supplementary figures 1,2,3). 3d models are provided in the GitHub repository.
  2. Using the 3D stereolithography files (*.stl) provided in the repository (https://github.com/BerezhnoyD/Reaching_Task_VAI) print all the required parts in PLA using a 3D-printer, including:
    - Lower and upper corners (4 for each) to hold the box together.
    - Two frames to attach mirrors to the box.
    - Base for the feeder motor and the feeder disk. **Note**: it is better to print the disk upside down to ensure it is smooth.
    - The XY-stage for easy and precise adjustment of the feeder position. **Note**
      i. We used the Ultimaker Cura Slicer and Ultimaker S3 3d printer. For most of the parts you should include support and an initial layer for better adhesion to the baseplate.
      ii. In case of any modifications needed, the original models from the Computer-aided design (CAD) software are also provided in the repository.
  3. Cut and drill 4 Plexiglas sheets (3 mm) for the side and back walls of the box according to the dimensions provided. The front wall should be made from thinner acrylic sheet (0.5mm). **Notes:**
    i. The front wall needs to be changed for each experiment, as it may get dirty and obstruct the view of the camera.
    ii. The frontal wall can be modified to have more room for head-mounted apparatus (e.g., miniscope, preamplifier).
    iii. CAD software along with CNC cutter machine can be used to scale and speed up the manufacturing process (all the design files provided were made using FreeCAD software, Open-source GNU license, https://www.freecad.org/), but the original design can be easily reproduced also with the use of all-manual tools.
  4. Assemble the box (Fig.1) using the Plexiglas sheets, corners, and screws (2mm) with nuts
    a. Nuts facing outside for easier maintenance.
    b. Screw the lower and upper corner in the proper places of the side walls and then attach the back wall to fit two parts together. The front wall should slide freely in place with no screws needed to allow for easy change.
    c. Screw the mirror frames into their places on both sides of the box and slide the mirrors in (should slide freely or the mirror frames will likely brake)
    d. Slide the metal rods in the holes at the bottom and glue them in place.
    **Notes:**
    i. Nuts should be facing outside for easier maintenance.
    ii. Grid floor makes the cleaning of the box easier and prevents animals from stashing the collected pellets in the box, which is important for successful training.
    iii. There is an elevated front rod, which is used as a starting point for animal to reach and a ‘perch’ where animals keep their paws in preparation for the reach. This rod should slide freely into its place and shouldn’t be glued for easier removal and cleaning.
    iv. The outside of the box can be covered with opaque film to provide dim-light condition.
  5. Solder 5-7 front floor rods together and attach them to a wire (15 cm) with a 2.54 mm pin connector on the other end (Fig.3C). Solder the elevated front rod to another wire with 2.54 mm pin connector. These two sets of rods are used as touch sensors needed to register the time spent near the slit, the beginning of the reaching trial etc.

**Fig.3.** Behavioral experiment controller based on the Arduino Nano board (A) and custom schematics on a breadboard enclosed in a box (Figure created with Fritzing software). The controller is connected to multiple electronic components of the behavioral box – automatic feeder (B) and floor bar sensors (C) with the use of a 10pin connector (A), which makes the assembly-disassembly of the box easy (D). The schematics are also provided separately on the Supplementary figure 4 and in the GitHub repository.
  6. Design of the motorized feeder. The motorized feeder is designed using a rotating disk to precisely position the pellet in front of the mouse without obscuring the view of the camera (Fig.1A, 3B). Assemble the 3D printed parts and attach the 28BYJ-48 stepper motor to the base/motor holder. The rotating disc plugs right into the shaft of the stepper motor and doesn’t require gluing. **Notes:**
    i. We designed two variants of the feeder – one for easier movement during the experiment (to dynamically adjust the distance from the slit) and the second one fixed and connected to the precise XY positioning stage. Designs for both are provided in the repository, but the first one is used in this protocol.
    ii. Using caution while positioning the stepper motor as a slight tilt or shift may affect the accuracy of the disc positioning. Check the precision of the pellet delivery before starting the real experiment.
  7. Positioning of the parts (Fig.1B).
    a. Assemble all the parts on a stable base. We used a thick piece of Plexiglas (5 mm) with drilled holes for wires and long screws. These upside-down screws, acting as anchor rods, make easier the precise positioning of the box every time by simply sliding screw shanks into the holes on the bottom angles. **Note**: Allow some space underneath or on the side for the wiring and the behavioral box controller.
    b. The same base is used to fix the XY positioning stage of the feeder and the video camera on a stable stand.
    c. The feeder should be positioned that way so the center of the closest pellet slot is 7 mm to the slit (5 mm from the edge of the disk to the slit). **CRITICAL:** The position and all settings of the video camera should remain the same throughout the whole experiment for the videos and 3D kinematics of the reaching to be comparable between days. The mirrors determining the angle of the side views are mounted to the box itself, so the parameters of the optical system are dependent mostly on the relative distance of the box to the camera. Thus, it is very important to fix the camera on the same stable surface as the behavioral box and the feeder, as well as fix the relative position of the latter to the camera.

### Design and assembly of the control schematics

#### Timing: 1 day

##### Design of the control schematics

A customized circuit is used to execute behavioral protocol, the stimuli and reinforcement (e.g., food pellets) presentation, data logging from the sensors, and video camera activation. The electrical components are housed in a small 3D printed box and soldered together on a breadboard (Fig.3D). The main component of the system is the Arduino Nano controller autonomously performing the programmed experimental protocol (written in C++ using Arduino IDE). Hence, this behavioral system can be used for (semi-)automatic training of mice to perform the reach-to-grasp task. Connected to a computer running custom Python script, this system collects both basic behavioral data from the touch sensors (position of the animal, paw placement, time spent in front of the slit, timing, and number of trials) and 3D view of the reaching movement (from the front camera and two side mirrors) for kinematic analysis.

1. Assembly of the breakout board. This is the interface between the components of the behavioral box. Position two Adafruit Industries AT424QT1070 capacitive touch sensor boards, ULN2003 Stepper Motor Drive, and IR Proximity Sensor on a breadboard and connect to the input and output 10 pin 2.54 mm sockets as pictured in Fig.3A. **Notes:**
  i. The power to the whole board is 5V delivered from the connection with Arduino board so all power and ground connections from all sensors should go to the single point at the 10-pin input (Arduino socket) – Power (red) and Ground (Brown) respectively.
  ii. The IR Proximity sensor serve as (1) an additional IR light source making the paw more contrast, and (2-optional) as a beam-break sensor detecting all the reaching attempts. For that purpose, we need to unsolder two LEDs from the sensor and mount them on the opposite ends of the slit facing each other: IR emitter at the bottom and detector on the top of the slit. They can be resoldered to the board with a long wire or use a pin-socket connection to the board.
  iii. All the other connections, both with Arduino and the behavioral box, are established using the 2.54 mm pin headers (See pinout on Fig.3A).
2. Solder the relay cables for the Arduino. There are three main relay cables: 10-pin connector for interface with the breadboard, 4-pin connector for controlling the feeder stepper motor and simple breadboard with two buttons to control the feeder rotation manually. All the schematics are provided in Fig.3. **Note:** We recommend mounting Arduino Board on the same breadboard or close to it while making the cable to the buttons longer and more durable.
3. Connect the control board to the behavioral box sensors (using the diagram on Fig.3A) and the video camera using Arduino pin 12 and GND and connecting them to the pin 1 - GPI and 6 – GND on the Blackfly S correspondingly (May be different for the camera you use). **Note**: Instructions on how to setup the synchronized recording on the FLIR Blackfly S camera used in this protocol can be found on FLIR official website (https://www.flir.com/support-center/iis/machine-vision/application-note/configuring-synchronized-capture-with-multiple-cameras/).

### Configuring computer for data streaming, storage, and analysis

#### Timing: 2-4 hours

1. Connect the Arduino Nano board to the computer and upload the experimental program. We have provided multiple programs written in C++ using the Arduino IDE (GPL), so the end user may either upload one of those or modify them to fit specific experimental needs. All of them are provided in the repository (https://github.com/BerezhnoyD/Reaching_Task_VAI; doi/10.5281/zenodo.7383917). The one used in the following protocol is the “Reaching_Task_Manual”.
  - ‘Reaching_Task_Draft’ – basic protocol with initialization for all the components to explore the structure of the program.
  - ‘Reaching_Task_Manual’ - Feeder is controlled manually and makes one step clockwise or counterclockwise when the experimenter presses the left or right button respectively.
  - ‘Reaching_Task_Feeder_Training’ – Feeder runs automatically and takes one step every 10sec while animal is in the front part of the box.
  - ‘Reaching_Task_Door_Training’ – If animal is detected in the front part of the box and grabs the elevated front rod, the feeder takes one step and the door blocking the slit opens (need additional servo motor connected for the door, see Fig.3A, socket pinout).
  - ‘Reaching_Task_CS_Training’ - If animal is detected in the front part of the box the speaker delivers 5s beep sound (CS trial) with intertrial intervals of 5s (ITI) and if the animal grabs the elevated front rod during this sound (trial) the feeder makes one step and the door blocking the slit opens (need additional servo motor and the speaker connected, see Fig.3A, socket pinout).
  **Notes:**
  i. After a single upload of the program the board will perform it autonomously on every power up. To switch to another protocol, you will need to connect Arduino to computer and upload another program using Arduino IDE.
  ii. The Arduino programming environment can be downloaded from the official website (https://www.arduino.cc/en/software) and is used to write, compile and upload the C++ code for Arduino controllers. It can also be used to stream the output of the behavioral box controller connected to the computer using the Serial Monitor (the proposed device outputs the data from all sensors as an updating table in a COM port interface).
2. Establish data streaming to PC. We use the Python scripts to record the data streamed from the FLIR camera and save it on the computer along with the sensor data from the behavioral box. That way the Arduino controls the experiment providing low latencies (10ms master clock) and precision needed for synchronization while PC handles only the visualization and saving of data for offline analysis. To start the experiment on the computer side you will need to run the script in Python. We have provided multiple example scripts that can be customized for your needs.
  - ‘CameraRoll.py’ – the script to run the FLIR camera in continuous recording mode, stream compressed video data to disk and save the data from Arduino sensors as a table (each column – one data stream) along with the corresponding FLIR camera frame number. All the adjustable parameters for the recording (ex. Exposure Time, Frame Rate, Time to record, Folder to Save Videos) are at the beginning of the Script. (**CRITICAL:** be sure to put in the right folder to save your videos to)
  - ‘CameraTrigger.py’ – the script to run the FLIR camera in triggered recording mode (triggered by the Arduino synchro pin), stream compressed video data to disk and display it on the screen + save the data from Arduino sensors as a table (each column – one data stream) along with the corresponding FLIR camera frame number. This is the Script used throughout the protocol as it saves only the important part of the video when the animal is holding the front bar and reaching continuously.
  **Notes:**
  i. You can locate these Scripts in the repository (ex. Reaching_Task_VAI/Recording toolbox/FLIR_GPU/CameraRoll.py) and copy them to the easily accessible folder on your PC.
  ii. As we use the Python scripts to interface with the FLIR camera we will need to install the FLIR Spinnaker and PySpin API from the official FLIR website (https://www.flir.com/products/spinnaker-sdk/) and Anaconda Python Platform (https://www.anaconda.com/products/distribution). Download and install the appropriate version of Spinnaker SDK from the FLIR website first. Then install Anaconda and check the Python version and only then install “Latest Python Spinnaker” checking it matches the installed version of Python you have.
  iii. The Python Script we use to handle the video still requires a few libraries. First, scikit-video (http://www.scikit-video.org/stable/) can be added to your Anaconda environment opening the Anaconda Prompt and typing in:

~~~
> pip install scikit-video
~~~ Second, FFMPEG - is mentioned in the script itself as a path to binary file. So the FFMPEG executable (https://ffmpeg.org/download.html) needs to be placed in a folder you can point to and then you should manually change the path in the Script accordingly. Opening the script <u>(ex. CameraRoll.py)</u> in Notepad look for the following line:

~~~
> skvideo.setFFmpegPath(‘C:/path_where_you_put_ffmpeg/bin/’)
~~~
  iv. Finally, you can plug both the Arduino board and FLIR Blackfly S camera to the PC USB ports and test the data acquisition. **CRITICAL**: Be sure to plug the video camera in the USB 3.0 port of the computer, otherwise you will experience a lot of dropped frames due to the USB interface speed limitations. There are two different Scripts provided. To run either of them open the Anaconda Prompt, start Python and point to one of the scripts to run it like in the example:

~~~
> python ‘C:/folder_with_a_script/CameraRoll.py’
~~~
3. Installation of the analysis software. All the scripts for general behavior and reaching movement kinematic analysis are written in Python and assembled in a series of Jupyter Notebooks. The user can perform step-by-step data analyses and visualization (Fig. 4). The scripts and the Notebooks can be downloaded from the project GitHub page (https://github.com/BerezhnoyD/Reaching_Task_VAI; <u>doi/10.5281/zenodo.7383917</u>), and to be able to run them you will need to install multiple Python libraries and dependencies. To make this process easier, we suggest running the installation through the Anaconda Python environment, handling all the dependencies properly.
  a. Download and install Anaconda Python Distribution through the official website (https://www.anaconda.com/)
  b. Download all the scripts and Notebooks from the project repository as well as DEEPLABCUT.yml file containing the all the dependencies needed to run the scripts (including the DeepLabCut and Anipose lib), which you will need to put in the folder accessible by Anaconda
  c. Install the environment using the Anaconda prompt command:

~~~
> conda env create -f DEEPLABCUT.yml
~~~
  d. If the environment setup was successful, you should have the new ‘DEEPLABCUT’ Anaconda environment which you should activate to run the analysis scripts. In Anaconda prompt run the following:

~~~
> cd path\to\the scripts\
> conda activate DEEPLABCUT
> jupyter notebook
~~~

**Fig.4.** Data processing workflow summarized in 3 Jupyter Notebooks run sequentially. Notebook 1 performs video tracking; Notebook 2 is reserved for manual behavioral analysis and labeling and Notebook 3 contains various data visualization scripts to generate the final plots. Figure was created with BioRender.com

This will open the Jupyter Notebook layout in your browser from which you will be able to navigate through the folder with the scripts and open the Notebook *.ipynb files with the main steps for analysis (Fig. 4).

## Key resources table

### Key resources table

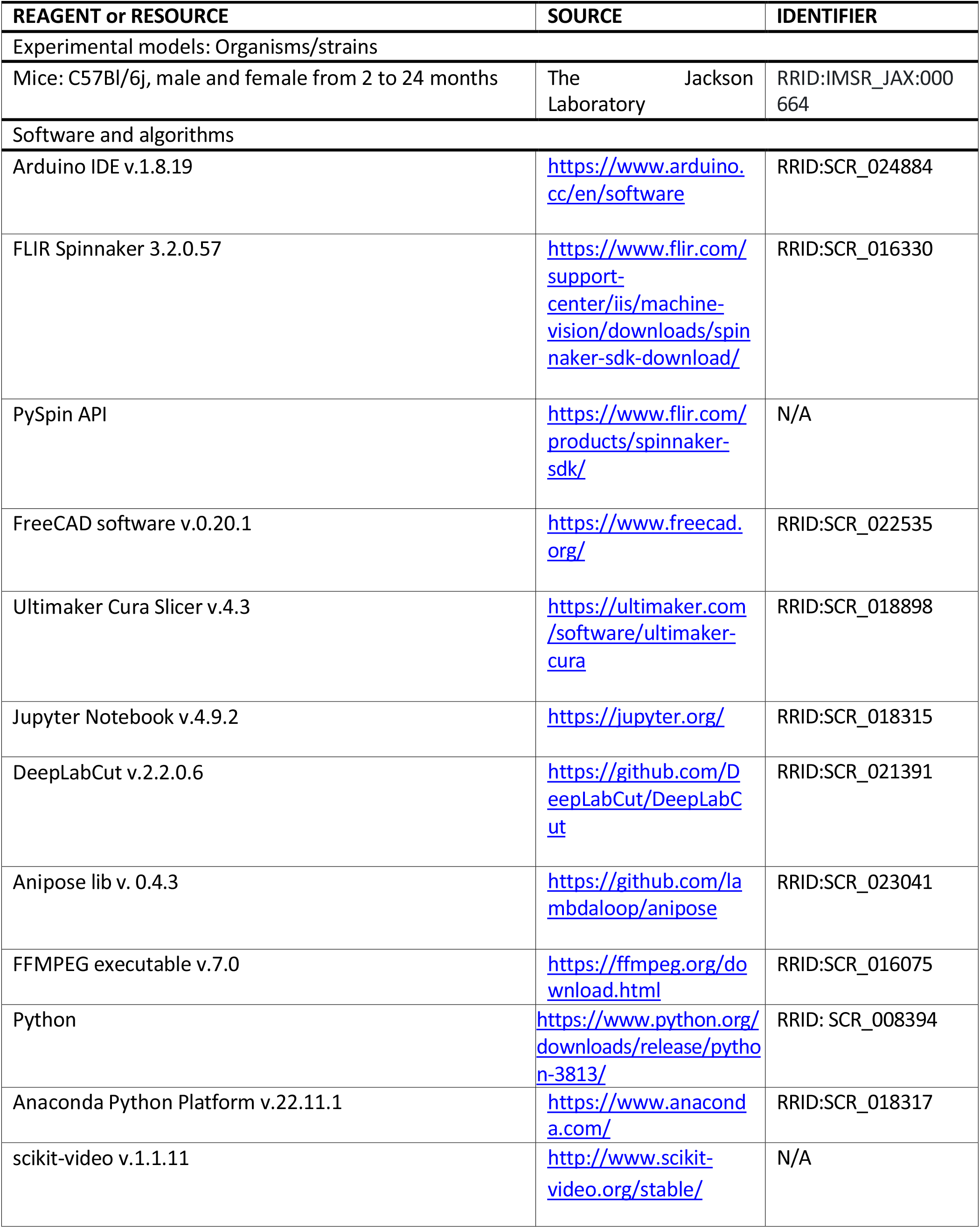
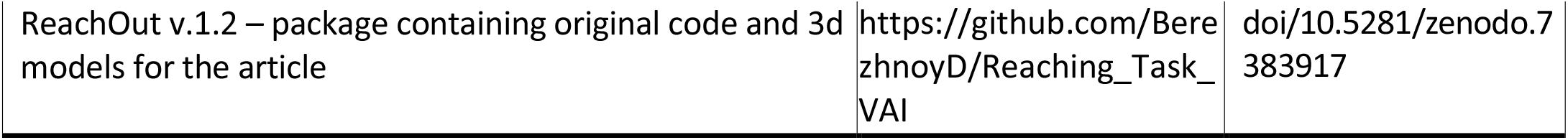
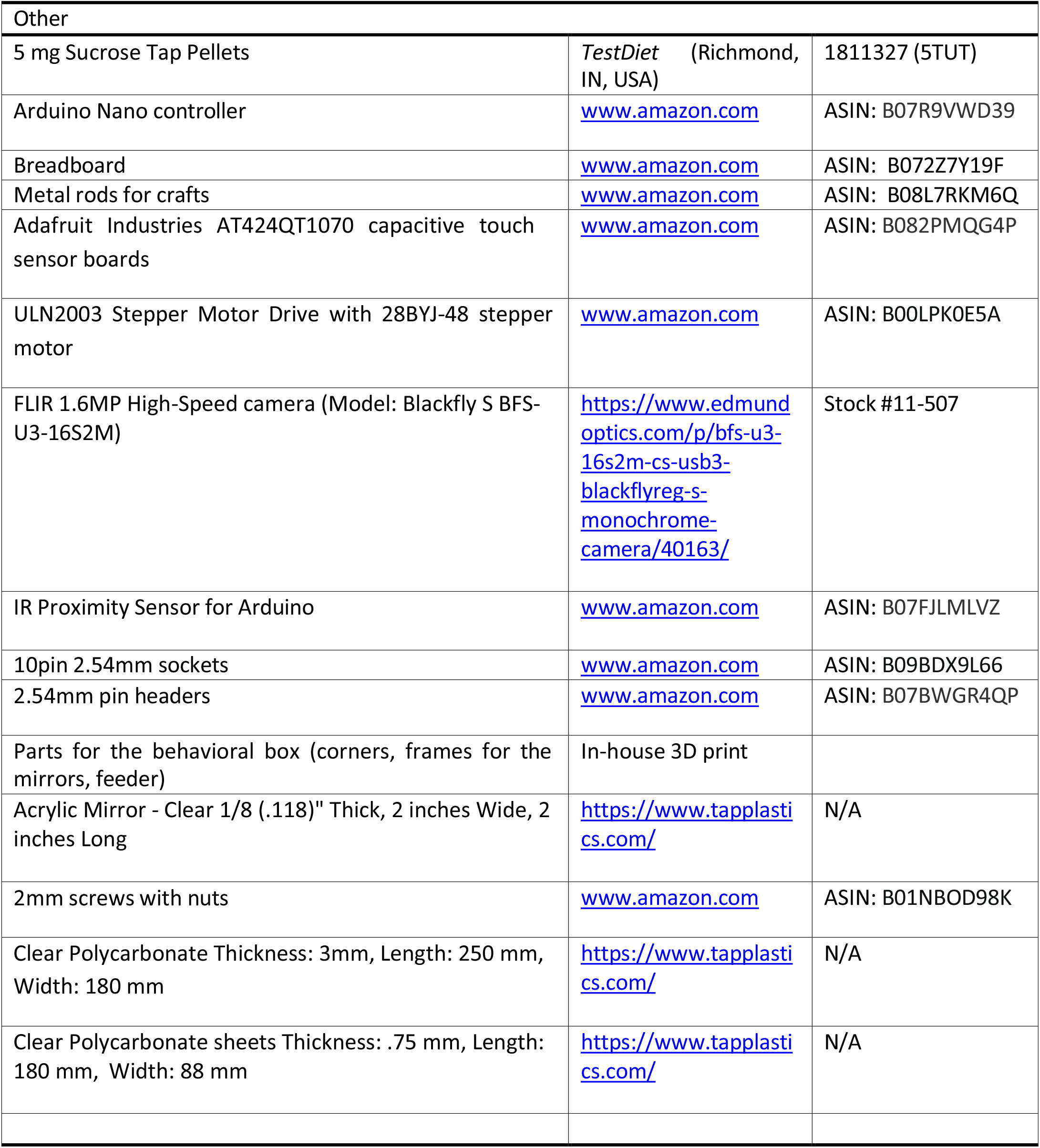

#### ▪ Mouse reach-to-grab task training protocol

The behavioral protocol consists of 5 days of habituation and 2 to 3 days of shaping followed by 7 days of training. Each of these stages will be detailed hereafter. All sessions are done during the light phase of the light/dark cycle.

**Note:** We noticed that mice are most motivated in the afternoon, when fed approx. 1 hour before the start of the dark phase of the light/dark cycle (16-hour food deprivation). Therefore, we planned all the experiments to end approx. 1 hour before the start of the dark phase. The sugar pellets used in the protocol were 5 mg spheres of approx. 2 mm in diameter (see. Key resources table).

#### Habituation

##### Timing: 5 days

###### Habituation to the experimenter (days 1-2)

We started each training session with 5 days of habituation that allows mice gradually habituate to the experimenter and the testing environment (e.g., room and behavior apparatus).

**Notes:**

i. During habituation, mice were food restricted. Throughout the experiment, make sure that the mice will be handled by the same experimenter until the end of training (e.g., for the weekly cage change).
ii. By the end of day 2 and to proceed to day 3, mice should get acclimated to the experimenter and handling (i.e., do not try to bite and circulate around experimenter’s hand in home cage, do not jump off experimenter’s hands/arms nor defecate and urinate excessively). If needed, extra days of habituation can be added, or the mice should be excluded from the study.

###### Habituation to the test setup I (day 3)

The purpose of this session is to acclimate the mice to the test box. To reduce stress, we propose that their first contact with the box be with a cage mate.

1. Transfer cage to the behavior room.
2. Allow mice to acclimate to the room for 5 minutes.
3. Place two mice from the same cage in the middle of the reaching box by holding tails.
4. Let the mice explore the box for 20 min.
5. Put animals back to their cage and add a few sugar pellets on the floor for consumption.
6. Clean the box with 70% ethanol between mice.

###### Habituation to the test setup II (days 4-5)

The purpose of the last two sessions of habituation is to allow animals get acclimated to the box individually, in the presence of sugar pellets. Recording the mouse behavior on habituation day 5 allows to assess this acclimation.

1. Transfer cage to the behavior room.
2. Allow mice to acclimate to the room for 5 minutes.
3. Place the feeder with a pile of sugar pellets 5 mm away from the front wall.
4. Place a plastic tray with 10 sugar pellets inside the box against the front wall. Note: no need to set up the elevated front rod at this stage.
5. Place a mouse in the center of the box.
6. Let the mouse explore the box for 20 min.
7. Move the mouse back to its cage.
8. Clean the box, the tray, and the disk with 70% ethanol between mice.

By the end of day 5, animals should show free exploration of the reaching box, interest in the slit and the pellets.

**Note:** Animals may not eat any pellets at this stage as they are not hungry but should be spending enough time near the slit and sniff it.

#### Shaping

##### Timing: 2-3 days

###### Overall procedure

Shaping stage is included to initiate reaching in simpler setting and determine the paw dominance before starting the actual training. Each session is recorded with the FLIR camera and the Arduino on, using the CameraRoll.py python script <u>in the PC.</u>

1. During the training animals should be food-restricted **Note**: In our lab, all animals were food-restricted throughout the shaping and training periods at the same level, i.e., around 80% of baseline bodyweight. The mice were housed in groups of 4 and food was placed on the cage floor. Daily food provided is the equivalent of 8% of animals’ baseline bodyweight.
2. Transfer the cage to the behavior room.
3. Allow mice to acclimate to the room for 5 minutes.
4. Place the feeder with a pile of sugar pellets 5 mm away from the front wall. **Note**. The disk can be placed closer to the slit and gradually moved away during the session. Also, the elevated front rod should not be set up during the shaping phase.
5. Connect the FLIR camera and the Arduino to a computer as previously described.
6. Place the mouse in the middle of the box.
7. Start the CameraRoll.py recording script to monitor the activity of the mouse for 20 min.
8. The mouse may retrieve sugar pellets by licking or reaching and the following numbers should be recorded:
  a. failed and successful licks
  b. failed and successful reaches with the right and left paws **Note**: When there are no sugar pellets remaining on the disk within the reaching distance from the slit, press the button to rotate the disk so that more sugar pellets are available to reach for.
9. Move the mouse back to the home cage at the end of the session.
10. Clean the box and the disk with 70% ethanol between mice.

For each mouse, calculate total number of reaches (**Equation 1**) and percentage of reaches with right paw (**Equation 2**) to determine paw dominance. The dominant paw is the paw used for more than 70% of all reaches (successful and failed).

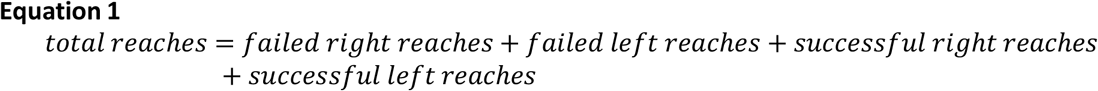

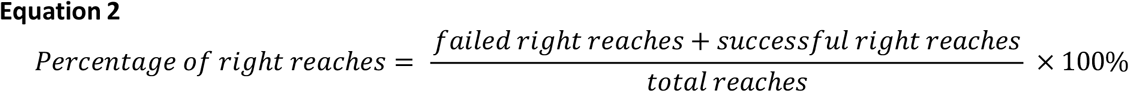

**Notes:**

a. Shaping stage takes 2 to 3 days. By the end of this stage, mice should be able to perform at least 20 reaches within 20 min and show paw dominance. Even if mice reach these criteria in shaping day 1, we strongly recommend keeping shaping day 2. Shaping day 3 is optional. If mice don’t meet this criterion, they are excluded from the study.
b. We highly recommend using the ‘Reaching_Task_Manual’ script during the Shaping phase and controlling the feeder with the buttons. The automatic scripts for the Arduino work well when the animal is reaching consistently.
c. Mice might start using their tongues to get sugar pellets (licking) before using their paws (reaching). In that case you can try moving the disc even further from the slit <u>(>9mm)</u>.
d. Shaping day 3 should be included if (i) the mouse still predominantly retrieves food pellets by licking at the end of shaping day 2, even if the mice have performed more than 20 reaches within the session, or (ii) if paw dominance can’t be determined at the end of shaping day 2.

#### Training

##### Timing: 7 days

###### Overall procedure

Training takes place after the shaping stage and requires at least 7 sessions (T1 to T7). Each session is recorded with the FLIR camera and the Arduino is on, using the CameraTrigger.py python script.

**Note**: Online observations are helpful to obtain preliminary results of each session (e.g., success rate and number of reaches), which can be used to optimize the training protocol whenever needed (see **Troubleshooting** below).

1. Transfer the cage to the behavior room.
2. Allow mice to acclimate to the room for 5 minutes.
3. Fill the slots of the feeder with sugar pellets.
4. Place the disk such as:
  a. Its edge is at 7 mm distance from the front wall.
  b. The sugar pellet is aligned with the left or right edge of the slit for mice showing right or left paw dominance, respectively.
5. Set up the elevated front rod.
6. Connect the FLIR camera and the Arduino box to the PC.
7. Place the mouse in the center of the box.
8. Start the CameraTrigger.py recording script (see **Optional**).
9. Monitor the activity of the mouse for 20 minutes.
10. Rotate the feeder using the buttons when the slot in front of the slit becomes empty.
11. Food pellets should be delivered:
  a. After successful reaches, when the mouse moves away from the slit to consume the pellet. **Note**: After food consumption, i f the mouse stays at the slit and keeps performing in vain reaches, no pellet should be delivered until the animal goes to the back of the reaching box and returns to the slit again.
  b. After failed reaches, when the animal goes to the back of the reaching box and comes to slit again.
12. At the end of the session, put the mice back to its home cage.
13. Clean the box and the disk with 70% ethanol between mice.
14. For each mouse calculate sum of failed and successful reaches and success rate (**Equation 3**).

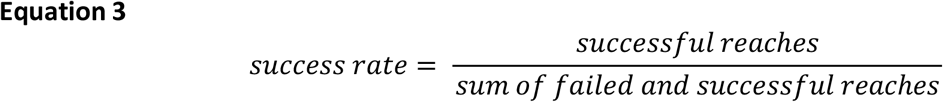

By the end of training, mice persistently reached for the pellets and conducted 100-300 reaches within 20 min. Around 70% of the animals trained using this protocol showed a success rate of 30-40% (Chen et al., 2014).

See **<u>Troubleshooting 1</u>**

###### Optional

If quantification of 3D trajectory (e.g., using Anipose) is needed, (1) the CameraRoll.py or CameraTrigger.py script should be started before each experiment; and (2) a 30 sec calibration video can be recorded for each day with a checkerboard calibration pattern visible in both central and mirror view. For more details on calibration see Anipose article^4^ or documentation in GitHub repository (https://github.com/lambdaloop/anipose).

#### Data analysis

##### Timing: 2-4 hours

###### Overall procedure

This part goes through the main processing steps of the analysis pipeline from opening the raw videos from the high-speed camera to the clustering and comparison of the 3d trajectories for different categories of reaches. The pipeline for the data analysis is shown on Fig.4

1. **ReachOut - Tracking**.**ipynb**: This notebook leads you through the main steps to convert the acquired video to the reconstruction of the mouse 3d paw trajectory. This Notebook relies on two state-of-the-art tools in markerless pose estimation and 3d triangulation: DeepLabCut^2^ and Anipose lib^4^.
  a. Single video acquired with the FLIR camera can be split into the left, central, and right views provided by the mirrors. **Note:** Both the behavior video and the corresponding calibration video from the same day should be split. Parameters of this cropping operation can be adjusted in the Script.
  b. Second cell in the Notebook is the command to start the DeepLabCut GUI interface, which can be used to open videos, label points for tracking, perform the training and evaluation of the neural network, and finally get the tracking for all the desired points (ex. snout, palm, 4 fingers, pellet). All the instructions for working with DeepLabCut can be found in the original repository (https://github.com/DeepLabCut/DeepLabCut). After running the DeepLabCut pipeline you should get the tracking for each point of interest in two projections (from central view and from the mirror view) to proceed with 3d reconstruction. Otherwise, you can use the 2d version of the tracking notebook provided in the repository as well. **Note:** To track the same points in two camera views we suggest running the DeepLabCut pipeline twice: once for the frontal view videos and the second time for the mirror (left or right depending on the dominant paw). We found that training two separate networks to track the same points in orthogonal camera views generates more accurate results than using one single network for all views. We suggest running the DeepLabCut pipeline twice: once for the frontal view videos and the second time for the mirror (left or right depending on the dominant paw).
  c. The last two steps in the processing notebook are dealing with the problem of paw triangulation using the Anipose Lib – restoring the coordinates in 3d space using two 2d trajectories from separate views.
    i. First, we will need to calibrate the camera – calculate the camera intrinsic and extrinsic parameters for our views. We use the calibration videos recorded right before each session with the small checkerboard pattern (4x4 squares, each square 1x1 mm), visible both from the frontal view and the mirror, and run the script on this video to acquire the calibration file. The script will automatically find the checkerboard pattern in the frames and ask you to confirm or reject the detection results manually (all points of the checkerboard should be detected and connected with lines from left to right, top to bottom) **Note:** This step could be done once for each batch of videos using single calibration video with a checkerboard for that day. Before running this script you will need to point the whole path to the calibration videos along with renaming them to match the Anipose pattern of A,B,C camera names. Further details can be found in the original Anipose repository (https://github.com/lambdaloop/anipose).
    ii. The previous step generates the calibration file (calibration.toml generated by Anipose) which we can apply to the 2d tracking data acquired with DeepLabCut for two different views (use the *.h5 files generated by DLC) to triangulate the points in 3d space. You will need to point the script to these 3 files and also point the path for the output file. **Note:** The names for the points to track should be the same in both DeepLabCut files. If you want to correct the coordinate frame to match certain static points in your video, you will also need to type in these points. They should also be present in the initial output files from DeepLabCut triangulation.
    iii. After this step you get the *.csv file with all the coordinates in absolute values (in mm, relative to the static points) and also can do a simple visual verification of the x,y,z coordinate for each of the part triangulated, which concludes the first Notebook.
2. **ReachOut - Analysis**.**ipynb**: This notebook contains the workflow to process the *.csv table with coordinates acquired on the previous step: clean the data, segment the trajectory to extract the relevant parts and assign the behavioral labels to the extracted parts. The screenshots for the following program snippets are shown in Fig.5

**Fig.5.** Processing the data and labeling the reaches using the proposed data analysis tools. The picture shows the GUI for different program snippets that the user runs through the processing pipeline (Notebook 2). A – extracting individual reaches from trajectory, B-looking at the videos for individual reaches and labeling the type of the reach. For the instructions on how to use the snippets read the annotated Notebook 2 in the GitHub repository.
  a. The first script (tracking_split) is designed to choose the parts of the trajectory for analysis (the peaks corresponding to the reach-to-grasp movement). It opens the *.csv file containing x,y,z coordinates for all the parts tracked and visualizes the trajectory for the selected part in 3d space (Fig.5A). The plot can be rotated and zoomed in/out by hovering on top of it with the mouse and holding the left mouse button. At first it shows the whole trajectory, but when you click on the progress bar in the bottom it will scroll through the small parts of the trajectory (500 frames at a time). Two smaller plots underneath the main one show the same trajectory projected on X (side view) and Y (front view) axis and are designed to extract even smaller parts of the trajectory - single reaches. When you click the left mouse button on these plots and move the cursor you will choose the part of the trajectory with a red span selector. If you want to save this part of the trajectory for further analysis, you should click the green “Save” button on the left. Scroll through the whole file, look at the trajectories and choose all the parts corresponding to the full reaches. When you finish analyzing the trajectory file you should click the green “Save_all” button on the right which saves the whole data frame with extracted parts of the trajectory for analysis as an *.h5 file.
  b. The second script (viewer) is opening the *.h5 file with the reaches extracted from the single video along with the video itself (*.mp4 file) and lets the user manually assign the category of the reach rewatching the video snippet corresponding to the trajectory extracted. The script opens the subplots with the selected trajectory from different views and two dropdown lists – one for the trajectories and one for the reach types (Fig.5B). You should sequentially choose each of the trajectories from the first list, which will open the corresponding video, and assign the category from the second list by simply clicking on it. The video can be closed by pressing the ‘Q’ button.
  c. By default, we classified the reaches to one of 6 categories depending on the trial outcome we have seen in the video:
    - Grasped – when mouse successfully grasped and consumed the food pellet.
    - Missed - when mouse did not touch the pellet during the reach.
    - Flicked – when mouse touched the pellet and knocked it down from the disk.
    - Lost – when mouse picked the pellet up but lost it on the way to the mouth.
    - In vain – animal reaching in the absence of food pellet.
    - Artifact – the recorded trajectory is not an acceptable reach. After accomplishing the classification, the script saves the *_scalar.h5 data frame with all the kinematic parameters for each of the reaches extracted. To open and visualize this data frame you should run the third analysis Notebook.
3. **ReachOut - Visualization**.**ipynb**: This notebook contains the scripts for visualization of the kinematic parameters, average projections, and additional automatic clustering of the extracted and labeled trajectories. The screenshots for the following program snippets are shown in Fig.6.

**Fig.6.** Visualizing the data using the proposed data analysis tools. The picture shows the GUI for different program snippets that the user runs through the processing pipeline (Notebook 3). A – mean trajectory visualization, B – scalar parameters visualization tool. For the instructions on how to use the snippets read the annotated Notebook 3 in the GitHub repository.
  a. The first script (reach_view) shows the average 3d trajectory along with projections of the reaching trajectories to 3 different axes to dissect the whole movement into its components: x (forward), y (sideward) and z (upward) (Fig. 6A). You can choose the category of reaches to show from the dropdown list and save the picture by clicking the right mouse button.
  b. The second script (scalar_view) plots all the kinematic parameters for the chosen category of reaches as the mean value (with mean parameter for individual reaches as the points) on the left and mean variance on the right (with the variation parameter, usually STD for individual reaches) (Fig. 6B). All the parameters plotted are calculated in the previous notebook and are taken from the *_scalar.h5 data frame. You should choose categories of reaches and parameters to show from the dropdown lists for the plots to be displayed.
  c. Third script (clustering) is optional and is designed to perform automatic clustering of the reaches based on the scalar parameters extracted. You should choose the clustering algorithm from the dropdown list and visualize the results. The results of the clustering can be saved to the *_scalar.h5 data frame as an additional labels column.
  d. The fourth script simply shows the number of reaches in each category labeled.
4. The third notebook concludes the analysis step and allows to generate the pictures reflecting main kinematic variables analyzed: reach trajectory, duration, velocity, reach endpoint coordinates etc.

## Expected outcomes

In this protocol we propose the open-source hardware-software solution for training mice to perform reach-to-grasp task, acquire the video for behavioral analysis and process it to analyze the fine motor skill kinematics. The proposed device can be easily manufactured in the lab with the readily available tools and materials with the only expensive component being the high-speed camera for behavioral acquisition. Behavioral apparatus can be used to perform multiple protocols depending on the study goals and not limited to the one suggested in the current study.

We used a high-speed camera with tilted mirrors to capture x, y and z coordinates of the movement in a single video. This approach acquires the data sufficient for 3D trajectory reconstruction using a single camera without synchronization of different video streams. Front- and side-view monitoring of the movement proved to be the most accurate in terms of paw and fingertip monitoring and sufficient to reliably distinguish between different movement outcomes. The analysis of the restored 3D trajectory provides accurate kinematic profile of the movement which can be further dissected into different directions (forward, sideward and upward, Fig. 7C) and phases (reaching, pronation, grasping, retraction, Fig. 5B-D) depending on the goal of the study. We demonstrate the effective manual and automatic clustering of the reaches into different categories and extraction of the number of kinematic parameters from every reach: endpoint coordinates, average and maximum velocity, acceleration and jerk, timing of the max velocity and peak positions and reach duration (Fig. 7A, B).

**Fig 7.** Exemplary data acquired with the system. A) Total number of reaches of wildtype C57BL/6 mice (n = 6) over the 7 days of training (T1 to T7). Results are presented as mean ± SEM. B) Success rate wildtype C57BL/6 mice (n = 6) over the 7 days of training (T1 to T7). The dotted line indicates the threshold (30%) chosen in the literature to consider a mouse as a “learner” mouse. C) Kinematic profile of the reach-to-grasp movement from a single animal acquired on the day 7 of training (T7) with the use of the proposed data analysis pipeline. Upper left plot shows all reaches from one representative mouse in 3D. All other panels show the dissection of the movement to three directional 2D components that can be analyzed separately – forward, sideward and upward motion of the paw. Data on each plot shows individual reaches in grey and averaged trajectory in red, Y-axis is expressed in distance to pellet.

Aside from movement kinematics which are analyzed from behavioral videos offline, experimenter may choose from several standard learning metrics like success rate or number of reaches in different categories to characterize the progress in a form of standard learning curves (Fig. 6A, B).

## Limitations

The current platform involves food deprived/restricted animals to perform fine movements with the forepaws, which could be an issue for animal models with motor impairments (e.g., Parkinson’s disease). Also, the success rate is highly dependent on the training protocol used and precise timing of the food delivery contingent upon animal actions. We found that extensive handling procedure prior to starting the training greatly improves the results, but also significantly increases the time required. In addition, only one animal can be present in the reaching box per training session. Thus, multiple units will be needed for high-throughput animal trainings.

In the data acquisition and analysis pipeline we used camera sampling rate 100 Hz, which along with lower resolution as a result of splitting the video to multiple views may cause occasional blurring during the fast parts of the movement, especially for the fingers. For studies focusing on individual digit control, increasing camera sampling rate (> 500 frame/sec) or using of multiple cameras will be needed to capture fine movement of fingers (Guo et al., 2015a; Becker and Person, 2019; Lopez-Huerta et al., 2021).

## Troubleshooting

1. It is quite often that animal training may not go as smoothly as described here or in literature. Mice might naturally lick off pellets instead of reaching for them, or not go to the back of the box or stay there. Some would simply exclude the mice from the protocol, but we chose to control the protocol manually and modify the protocol based on observed behaviors.
2. Training environment is key to a successful training:
  - Behavioral room should be quite with proper temperature and lighting.
  - There should be only the one experimenter in the room. The experimenter shouldn’t wear any kind of perfume.
  - If a mouse jumps out of the hand during habituation to the experimenter, it can be returned to the home cage. The habituation can be tired again after habituating the rest of the mice.
3. Tips to engage animals into training”
  - On shaping day 1, if the mouse shows no interest in getting food (no licks, no reaches), you can try to provide sugar pellets in the home cage with regular chow at the end of day, but still maintain the same total amount of food provided (e.g. 2g of chow + 2g of sugar pellets can be provided for calculated 4 g food daily. Make sure to add an additional shaping day.
  - During the initial days of training, if the mouse starts biting on the slit or performs more than 20 consecutive in-vain reaches, a brief noise can be introduced by gently scratching the top corner of the side wall. This will distract the mouse and move it to the rear of the reaching box. We have noticed that the mouse would then come forward again after hearing the rotating disk.
  - If the mouse stays in the back of the test box for more than 5 seconds, rotate the disk again. The sound of the rotating disk might encourage the mouse to come forward.
  - At the beginning of single-pellet training, if the mouse sniffs through the slit but does not reach, adding a small pile of pellets in the center of the disk encourages the mouse to reach.
4. Tips to encourage reaching and prevent licking:
  - If the mouse attempts to lick off the pellets during the early days of training, move the disk slightly away from the slit. This will prevent the pellet from being reachable by tongue but still reachable by paw. In case the mouse stops licking for a full session, move the disk closer to the slit in the next session. Move 1 to 2 mm closer every 5 minutes, as long as the mouse keeps reaching, until you can place it at the standard 5 mm distance from the front wall.
  - If the mouse loses too much weight (i.e. < 75% of baseline body weight), it has a tendency to start licking, even after the shaping phase. If the previous solution to licking does not work, we found it helpful to increase the food portion after the session and move back to the shaping setup on the next session.
5. Tips to increase success rate:
  - We strongly encourage a single experimenter to conduct the habituation and behavioral shaping and training sessions. We noted that animals can stop or reduce reaching when a new experimenter is involved at any stages of the task. For instance, even well trained (i.e., “experts”) mice could stop reaching when additional experimenter assisted with the training.
  - Shaping and training sessions are expected to be performed at the same time of the day to reduce variabilities due to circadian rhythm.
  - Successful acquisition of this reaching skills requires daily training for at least consecutive 4 days after the completion of shaping.
  - We were using a rotating disk for pellet delivery. Some modifications may be considered and can be helpful to increase success rate of reaches, such as slightly increasing depth of wells, adjusting the height of disk, and precisely positioning the food dispenser.

## Resource availability

### Lead contact

Further information and requests for resources and reagents should be directed to and will be fulfilled by the lead contact, Hong-yuan Chu.

### Materials availability

This study did not generate new unique reagents. All the devices listed in this study can be either found in key resources table or manufactured in-lab.

### Data and code availability

The datasets and code used during this study are available at https://github.com/BerezhnoyD/Reaching_Task_VAI

## Acknowledgments

The authors thank Van Andel Institute Maintenance Department for the assistance in customizing the mouse reach training box and Van Andel Institute Research Operation team for 3D printer and supplies. This research was funded in whole or in part by Aligning Science Across Parkinson’s (ASAP-020572) through the Michael J. Fox Foundation for Parkinson’s Research (MJFF). This work was partially supported by National Institute of Neurological Disorders and Stroke grant: R01NS121371 (H.-Y.C.). For the purpose of open access, the authors have applied a CC BY public copyright license to all Author Accepted Manuscripts arising from this submission.

## Author contributions

DaniilBerezhnoi: Methodology, Software, Visualization, Writing-Original Draft, Editing

Hiba Douja Chehade: Methodology Validation, Investigation, Visualization, Writing-Original Draft, Editing

Hong-Yuan Chu: Funding acquisition, Supervision, Conceptualization, Resources, Visualization, Writing-Review & Editing

## Figure legends

Supplementary Fig.1 The blueprint showing all the dimensions for the sidewall of the box. The CAD model can be found in the repository.

Supplementary Fig.2 The blueprint showing all the dimensions for the frontal wall of the box. The CAD model can be found in the repository.

Supplementary Fig.3 The blueprint showing all the dimensions for the back wall of the box. The CAD model can be found in the repository.

Supplementary Fig.4 The blueprint showing the enlarged circuitry and positioning of the components on the breadboard. The Fritzing file can be found in the repository.

## CELL PRESS DECLARATION OF INTERESTS POLICY

Transparency is essential for a reader’s trust in the scientific process and for the credibility of published articles. At Cell Press, we feel that disclosure of competing interests is a critical aspect of transparency. Therefore, we require a “declaration of interests” section in which all authors disclose any financial or other interests related to the submitted work that (1) could affect or have the perception of affecting the author’s objectivity or (2) could influence or have the perception of influencing the content of the article.

### What types of articles does this apply to?

We require that you disclose competing interests for all submitted content by completing and submitting the form below. We also require that you include a “declaration of interests” section in the text of all articles even if there are no interests to declare.

### What should I disclose?

We require that you and all authors disclose any personal financial interests (e.g., stocks or shares in companies with interests related to the submitted work or consulting fees from companies that could have interests related to the work), professional affiliations, advisory positions, board memberships (including membership on a journal’s advisory board when publishing in that journal), or patent holdings that are related to the subject matter of the contribution. As a guideline, you need to declare an interest for (1) any affiliation associated with a payment or financial benefit exceeding $10,000 p.a. or 5% ownership of a company or (2) research funding by a company with related interests. You do not need to disclose diversified mutual funds, 401ks, or investment trusts.

Authors should also disclose relevant financial interests of immediate family members. Cell Press uses the Public Health Service definition of “immediate family member,” which includes spouse and dependent children.

### Where do I declare competing interests?

Competing interests should be disclosed on this form as well as in a “declaration of interests” section in the manuscript. This section should include financial or other competing interests as well as affiliations that are not included in the author list. Examples of “declaration of interests” language include:

> “AUTHOR is an employee and shareholder of COMPANY.”
>
> “AUTHOR is a founder of COMPANY and a member of its scientific advisory board.”

*NOTE*: Primary affiliations should be included with the author list and do not need to be included in the “declaration of interests” section. Funding sources should be included in the “acknowledgments” section and also do not need to be included in the “declaration of interests” section. (A small number of front-matter article types do not include an “acknowledgments” section. For these articles, reporting of funding sources is not required.)

### What if there are no competing interests to declare?

If you have no competing interests to declare, please note that in the “declaration of interests” section with the following wording:

> “The authors declare no competing interests.”

## CELL PRESS DECLARATION OF INTERESTS FORM

If submitting materials via Editorial Manager, please complete this form and upload with your initial submission. Otherwise, please email as an attachment to the editor handling your manuscript.

***Please complete each section of the form and insert any necessary “declaration of interests” statement in the text box at the end of the form. A matching statement should be included in a “declaration of interests” section in the manuscript***.

### Institutional affiliations

We require that you list the current institutional affiliations of all authors, including academic, corporate, and industrial, on the title page of the manuscript. ***Please select one of the following:***

☒ All affiliations are listed on the title page of the manuscript.
□ I or other authors have additional affiliations that we have noted in the “declaration of interests” section of the manuscript and on this form below.

### Competing financial interests

We require that authors disclose any financial interests and any such interests of immediate family members, including financial holdings, professional affiliations, advisory positions, board memberships, receipt of consulting fees, etc., that:

1. could affect or have the perception of affecting the author’s objectivity, *or*
2. could influence or have the perception of influencing the content of the article.

### Please select one of the following

☒ We, the authors and our immediate family members, have no financial interests to declare.
☒ We, the authors, have noted any financial interests in the “declaration of interests” section of the manuscript and on this form below, and we have noted interests of our immediate family members.

### Advisory/management and consulting positions

We require that authors disclose any position, be it a member of a board or advisory committee or a paid consultant, that they have been involved with that is related to this study. We also require that members of our journal advisory boards disclose their position when publishing in that journal. ***Please select one of the following:***

☒ We, the authors and our immediate family members, have no positions to declare and are not members of the journal’s advisory board.
□ The authors and/or their immediate family members have management/advisory or consulting relationships noted in the “declaration of interests” section of the manuscript and on this form below.

### Patents

We require that you disclose any patents related to this work by any of the authors or their institutions.

### Please select one of the following

☒ We, the authors and our immediate family members, have no related patents to declare.
□ We, the authors, have a patent related to this work, which is noted in the “declaration of interests” section of the manuscript and on this form below, and we have noted the patents of immediate family members.

***Please insert any “declaration of interests” statements in this space***. This exact text should also be included in the “declaration of interests” section of the manuscript. If no authors have a competing interest, please insert the text, “The authors declare no competing interests.”

The authors declare no competing interests

☒ **On behalf of all authors, I declare that I have disclosed all competing interests related to this work. If any exist, they have been included in the “declaration of interests” section of the manuscript**.

