## Supplementary figures and images for "Open-Source Platform for Kinematic Analysis of Mouse Forelimb Movement" (from another version of this preprint: version 2, 2024-04-15)

### Supplement figure 1

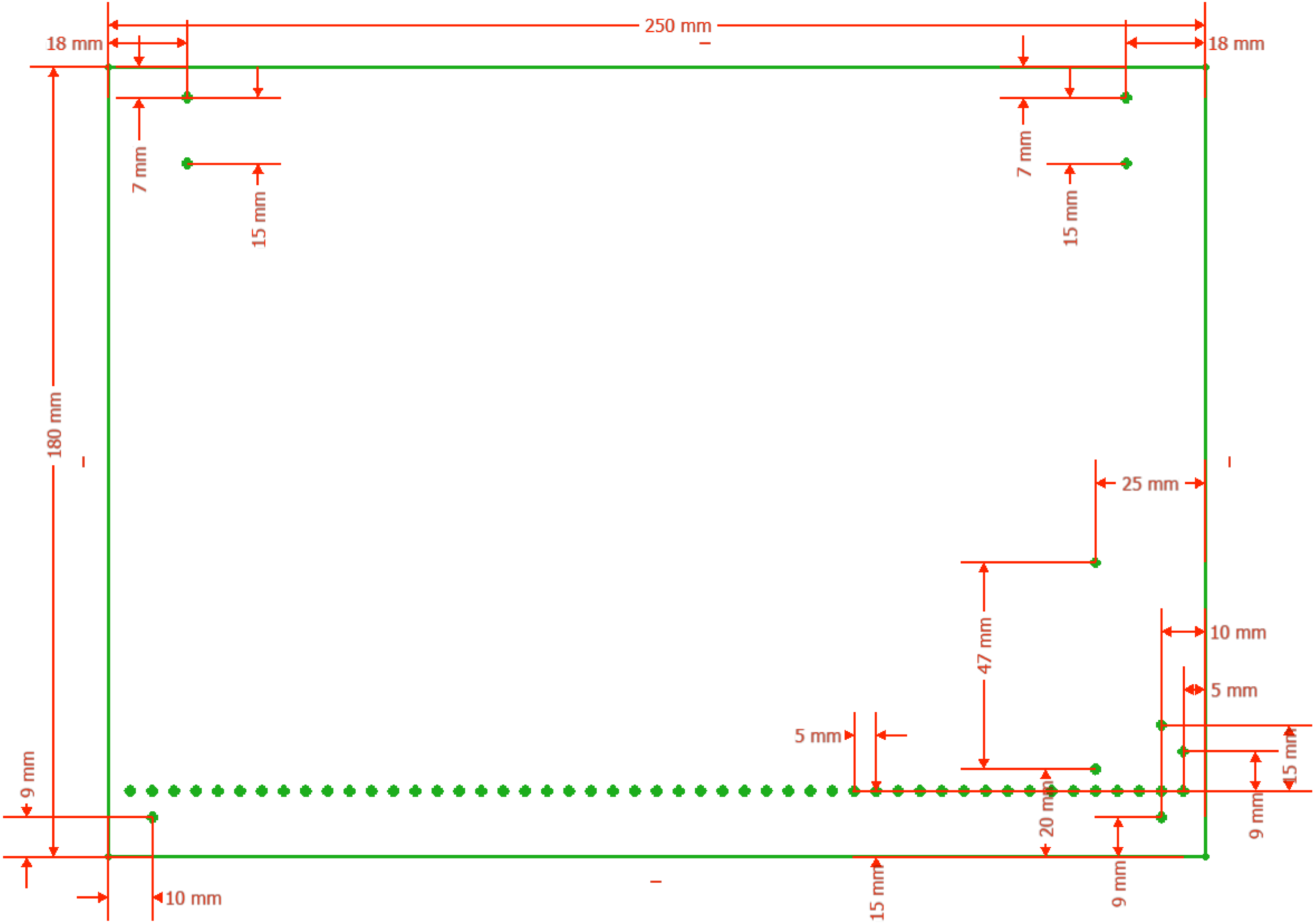

### Supplement figure 2

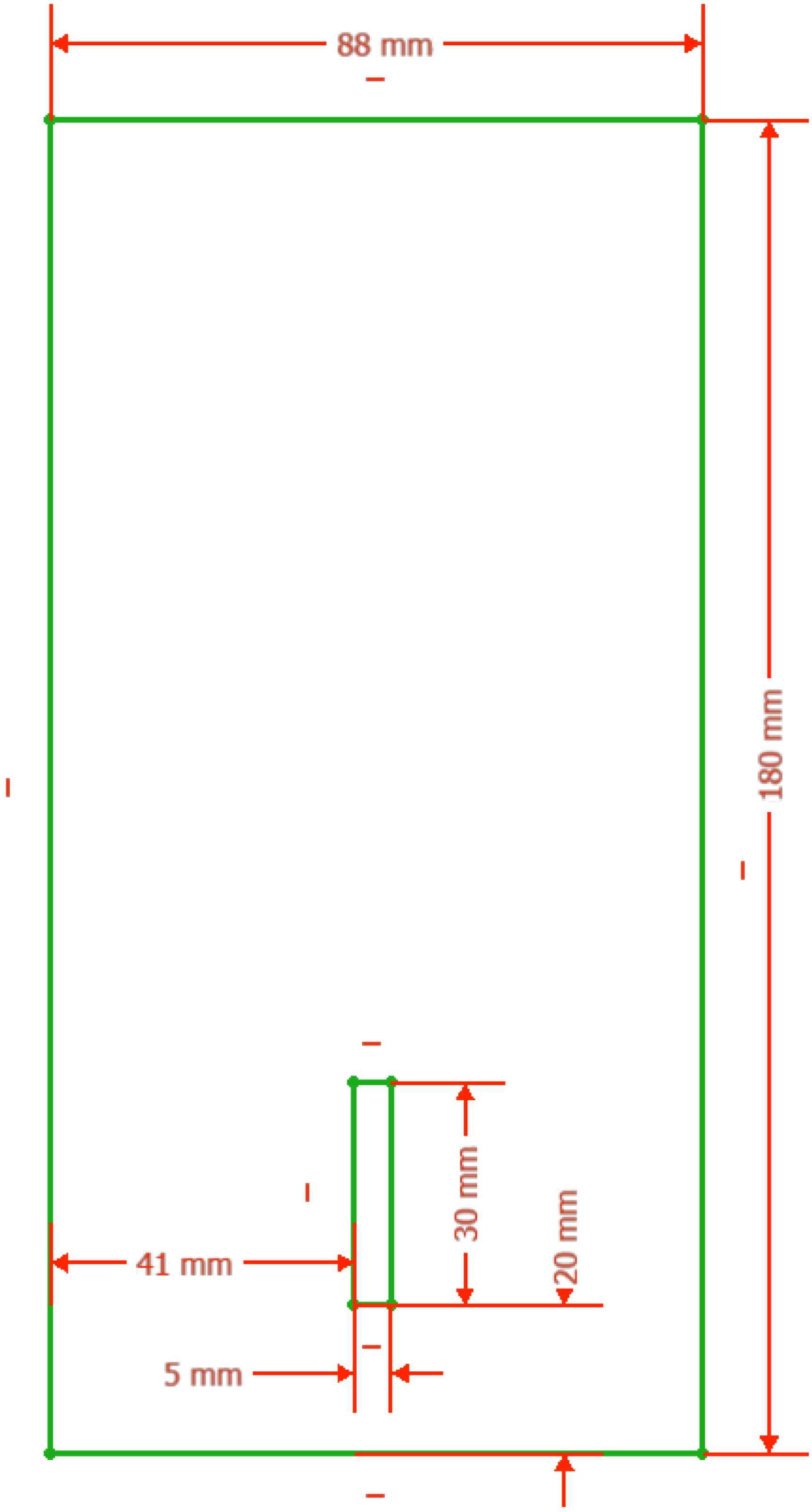

### Supplement figure 3

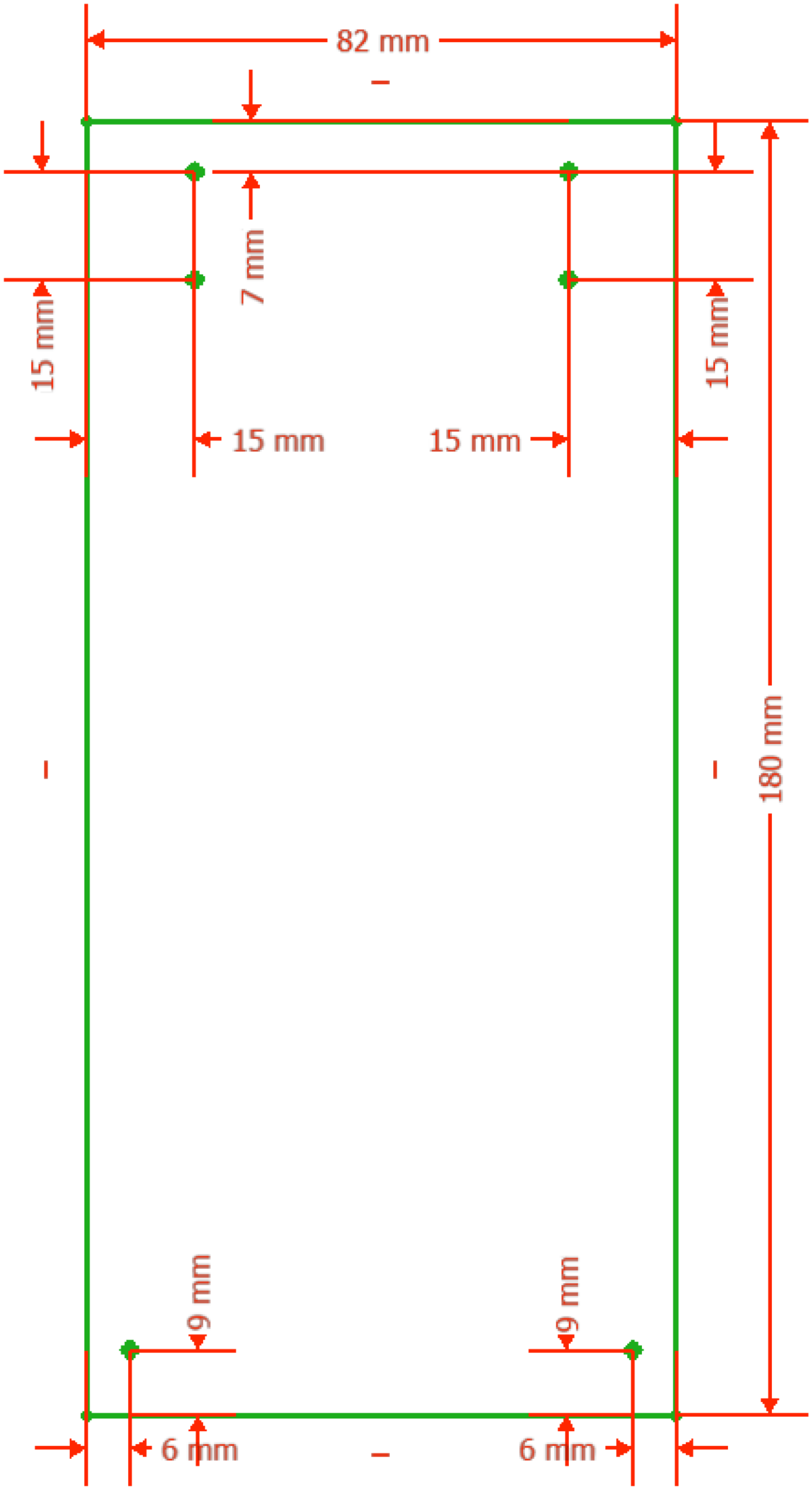
