## Supplement figure 4 for "Open-Source Platform for Kinematic Analysis of Mouse Forelimb Movement" (from another version of this preprint: version 2, 2024-04-15)

**Main components:**

1 – control buttons
2 – 4pin connector to 4
3 – 10pin connector to 6
4 – 4pin socket
5 – stepper motor socket to 8
6 – 10pin Arduino socket
7 – 10pin output socket
8 – feeder stepper motor
9 – beam sensor
10 – touch sensors

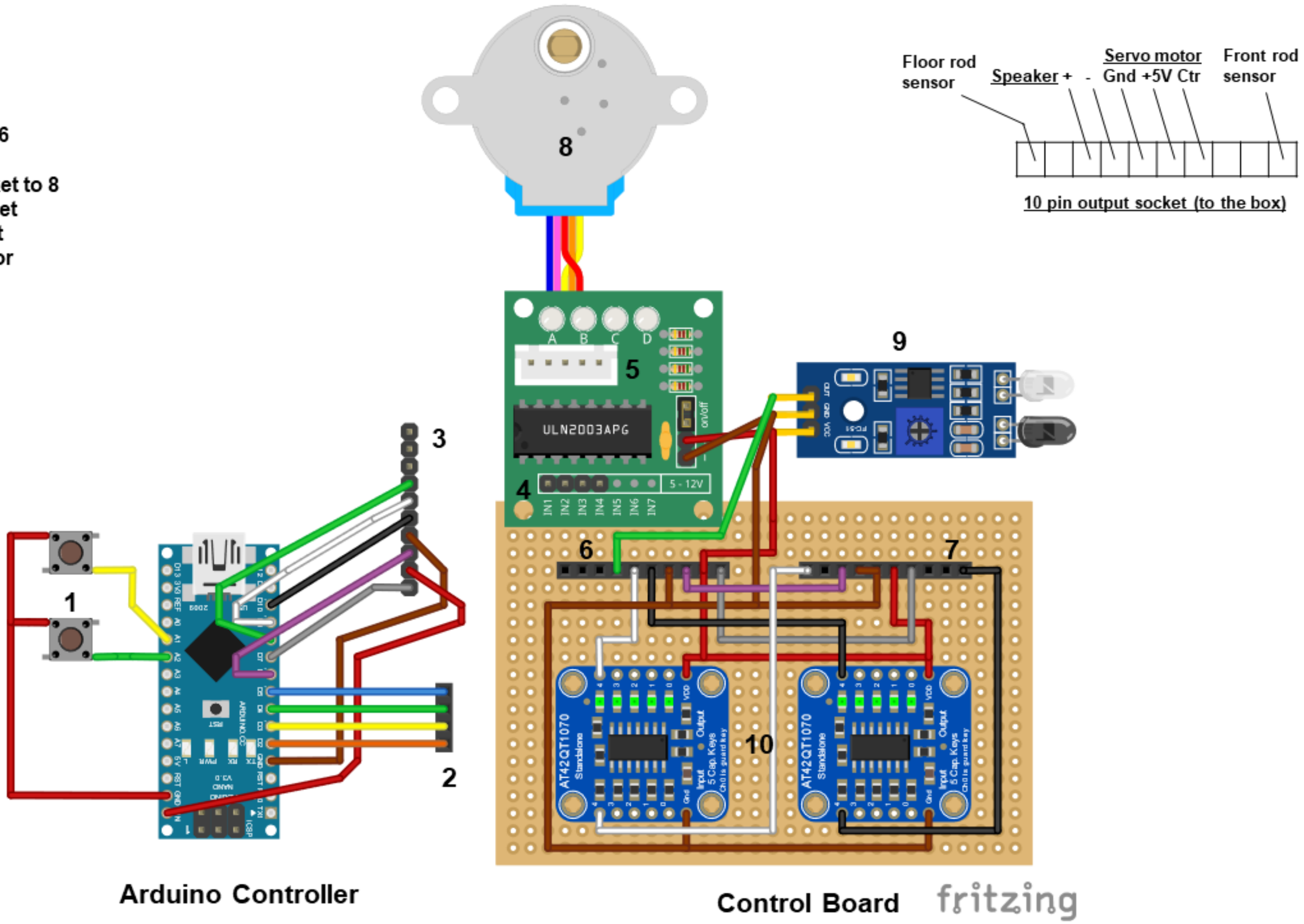
